# DLL4 and VCAM1 enhance the emergence of T cell-competent hematopoietic progenitors from human pluripotent stem cells

**DOI:** 10.1101/2021.11.26.470145

**Authors:** Yale S. Michaels, John M. Edgar, Matthew C. Major, Elizabeth L. Castle, Carla Zimmerman, Ting Yin, Andrew Hagner, Charles Lau, M. Iliana Ibañez-Rios, David J.H.F. Knapp, Peter W. Zandstra

## Abstract

T cells are key mediators of the adaptive immune response and show tremendous efficacy as cellular therapeutics. However, obtaining primary T cells from human donors is expensive and variable. Pluripotent stem cells (PSCs) have the potential to serve as a consistent and renewable source of T cells, but differentiating PSCs into hematopoietic progenitors with T cell potential remains a significant challenge. Here, we developed an efficient serum- and feeder-free protocol for differentiating human PSCs into hematopoietic progenitors and T cells. This defined method allowed us to study the impact of individual recombinant proteins on blood emergence and lineage potential. We demonstrate that the presence of DLL4 and VCAM1 during the endothelial-to-hematopoietic transition (EHT) enhances downstream progenitor T cell output by >80-fold. Using single cell transcriptomics, we showed that these two proteins synergise to drive strong notch signalling in nascent hematopoietic stem and progenitor cells and that VCAM1 additionally drives a pro-inflammatory transcriptional program. Finally, we applied this differentiation method to study the impact of cytokine concentration dynamics on T cell maturation. We established optimised media formulations that enabled efficient and chemically defined differentiation of CD8αβ+, CD4-, CD3+, TCRαβ+ T cells from PSCs.

## Introduction

T cells are potent therapeutic agents. Chimeric antigen receptor (CAR) and engineered TCR-T cells have demonstrated robust clinical efficacy against cancer^1–3^ and show promise for treating infection and immunological disorders as well as preventing transplant rejection^4, 5^. However, sourcing T cells from individual donors is expensive, time consuming and laborious ^6^. Inter-donor variability and intra-donor heterogeneity in CAR/TCR transduction make personalised T cell therapies difficult to manufacture and quality control^7–9^.

Human pluripotent stem cells (hPSC) offer an attractive solution to these manufacturing challenges^10^. hPSC have the capacity for unlimited self-renewal. They can be engineered, clonally selected and screened for quality control prior to expansion, enabling scalable production of a homogenous product^11^. Developing efficient processes for differentiating hPSCs into T cells will enable cost-effective and consistent cell therapy manufacturing, ultimately leading to more efficacious products^12–17^.

A major bottleneck on the path to T cell production is to efficiently differentiate hPSCs into definitive hematopoietic progenitors with T cell potential^18–21^. Definitive hematopoietic stem cells (HSCs) arise from a cell type known as hemogenic endothelium in a process called the endothelial-to-hematopoietic transition (EHT)^22, 23^. Previous methods for producing T-lineage competent hematopoietic stem/progenitor cells from hPSCs have used immortalised stromal cell lines and serum or undefined cellular extracts to support EHT and T cell differentiation^12, 14, 15, 17, 20, 24^. These undefined components obfuscate the signalling interactions that drive development, making key parameters difficult to identify and optimise. The use of animal-derived sera, feeder cells and undefined extracts also causes lot-to-lot variability and limits translation to the clinic^25^.

Iriguchi et al. recently developed a feeder-free protocol for differentiating iPSCs into T cells^13^. Using iPSCs that already harboured re-arranged TCRs, either by reprogramming from a T cell clone or through TCR transduction, authors were able to generate CD8+, CD4-, CD3+, TCRαβ+ T cells. Despite this important advance, iPSCs without a pre-rearranged TCR failed to develop into TCRαβ+ T cells, suggesting their process failed to capture key aspects of conventional T cell development^13^.

In another recent study, Trotman-Grant et. al. succeeded in generating CD4+, CD8+, CD3+, TCRαβ+ T cell progenitors from iPSCs without the need to supply a re-arranged TCR, achieving an important milestone^16^. But in this protocol, the efficiency of converting CD34+ cells to T cell progenitors cells was low (estimated by the authors to be on the order of 1/1000), consistent with a bottleneck at the stage of generating lymphoid-competent hematopoietic cells^16^. The authors first generated CD34+ cells in a mesoderm/hematopoietic differentiation, which they subsequently seeded into a T cell differentiation culture^16^. The protocol that they used to generate CD34+ cells has been reported to primarily produce hemogenic endothelial cells, the developmental pre-cursor of HSC^21^. These cells may not be at the appropriate developmental stage to begin T cell differentiation.

Here, we hypothesized that adding an EHT stage to the differentiation process would allow us to convert cells to lymphoid competent hematopoietic progenitors at the appropriate developmental stage to progress into T cell differentiation conditions. We aimed to use recombinant proteins, rather than stromal cell lines, serum or undefined extracts, to enable better control over the EHT process and to gain insight into the biological requirements for lymphoid-competent hematopoiesis.

Prior work suggests that notch signalling during EHT is required to produce lymphoid competent hematopoietic cells^24, 26^. This motivated us to test the potent notch ligand DLL4 in our culture system. Transcriptional profiling in mouse revealed that the cell adhesion molecule VCAM1 is highly enriched in cells of aorta-gonad-mesonephros (AGM) capable of supporting definitive hematopoiesis^27^. Our lab has shown that VCAM1 is able to synergise with DLL4 to enhance notch signalling in a different developmental context^28^. Consequently, in this study we also tested the impact of VCAM1 during EHT. We show that DLL4 and VCAM1 interact to drastically improve production of hematopoietic progenitors with T cell potential from hPSCs. Collectively, the addition of these two proteins during EHT increased downstream T cell progenitor output by more than 80-fold.

The chemically defined differentiation method we developed here allowed us to probe the impact of each immobilised protein on hematopoietic lineage output and transcriptional programs. This analysis revealed that VCAM1 promotes an inflammatory transcriptional program in emergent hematopoietic stem and progenitor cells. Finally, this method enabled the production of not only T cell progenitors, but mature CD8αβ+, CD4-, CD3+, CD62L+, TCRαβ+ T cells and allowed us to study the impact of cytokine exposure dynamics on T cell maturation. Our platform provides a customisable environment capable of revealing key signalling requirements for blood emergence and T cell development. This new method will contribute to robust, clinically compatible manufacturing of PSC-derived T cell therapies.

## Results

### DLL4 and VCAM1 enhance the emergence of T cell competent hematopoietic progenitors from human pluripotent stem cells

We set out to understand the minimal signalling requirements for generating hematopoietic progenitors with T cell potential from hPSCs. We first attempted to generate hemogenic endothelial (HE) cells, the developmental precursors of hematopoietic stem cells, by aggregating hPSC into 3D structures by centrifugation in microwell plates^29^. We subjected the aggregates to a step-wise series of media formulations known to specify mesoderm, and subsequently definitive haemato-endothelial identity^21, 26^. We dissociated the resulting aggregates and separated CD34+ cells to enrich for HE. We placed the CD34+ cells onto an uncoated tissue culture surface in media formulated to promote endothelial-to-hematopoietic transition (EHT)^30^ (Figure 1a). We observed a morphological transition from adherent, endothelial-like cells, to non-adherent spherical cells (Supplementary Figure 1).

**Figure 1:**
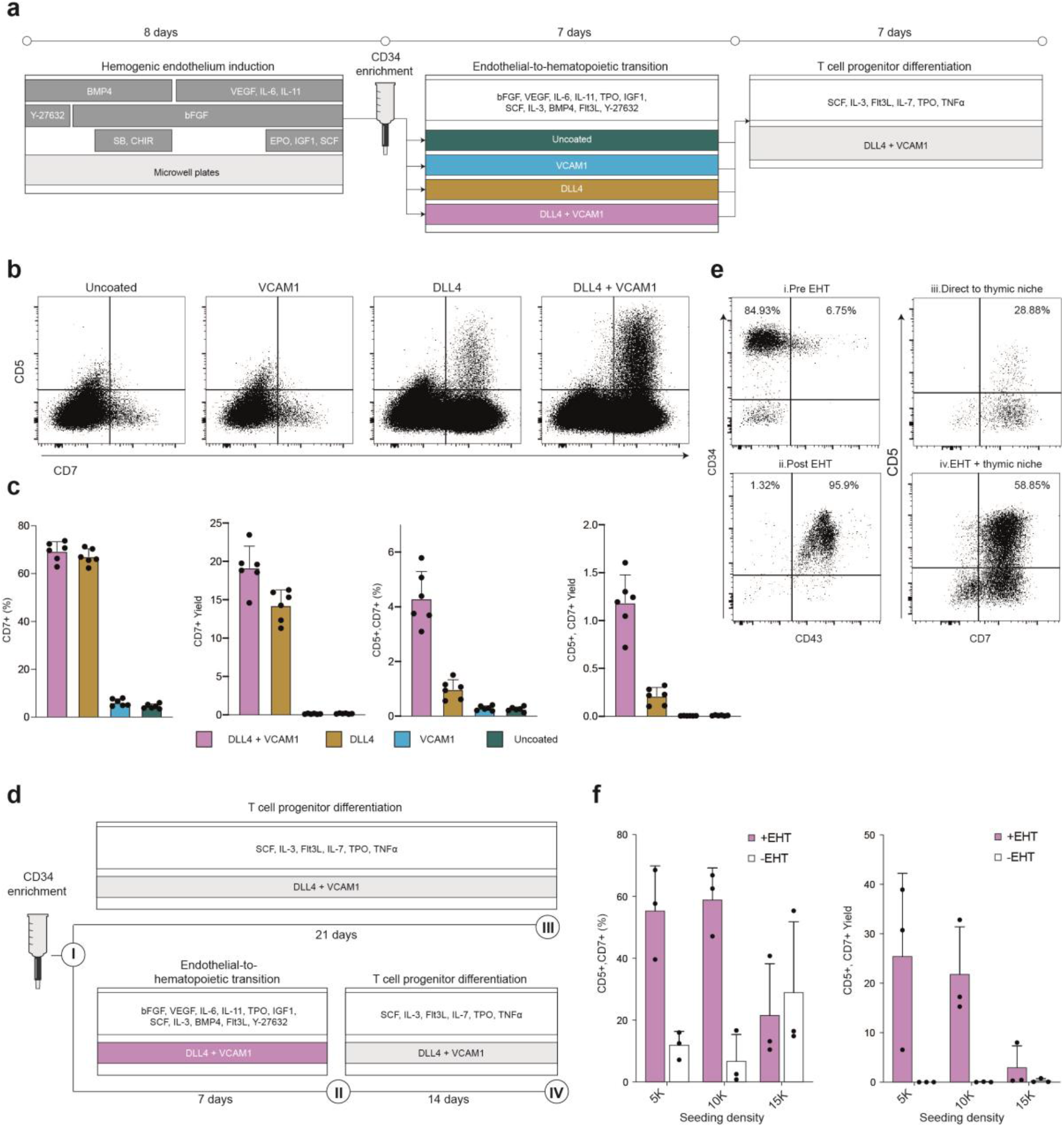
The presence of DLL4 and VCAM1 during the endothelial to hematopoietic transition supports development of HSPC with robust T cell potential. **a.)** Schematic overview of chemically defined platform for producing multipotent hematopoietic progenitors and T cell progenitors from pluripotent stem cells. **b.)** Flow cytometry analysis of progenitor T cell output after transitioning cells from 7 days in each EHT coating condition into a common defined thymic niche for an additional 7 days. **c.)** Quantification of the frequency and yield of CD7+ lymphoid progenitors and CD7+, CD5+ T cell progenitors after 7 days in the thymic niche (mean +/- s.d., n=6). **d.)** Experimental design to assess the effect of adding or omitting the EHT culture phase prior to transferring cells into the thymic niche. e.) Immunophenotype of cells generated with or without the EHT culture stage. Numerals in **(e)** correspond to the schematic in **(d). f.)** Quantification of the yield and frequency of T cell progenitors generated with or without an EHT culture phase.

Next, we assayed whether these non-adherent hematopoietic cells were capable of differentiating into T cell progenitors. We transferred the cells to plates coated with immobilized DLL4, a notch ligand capable of promoting T lineage specification, and VCAM1, a cell adhesion molecule that we have previously shown to enhance this process^28^ (Figure 1a). Though we cultured the cells in media capable of generating CD5+, CD7+ T cell progenitors from cord-blood stem cells, we detected almost no CD5+, CD7+ cells from hPSC derived hematopoietic cells (Figure 1b,c). In fact, we observed very low numbers of CD7+ cells, an earlier and less committed progenitor (Figure 1b,c).

Multiple reports suggest that notch signalling during EHT promotes the emergence of HSPC with T lineage potential^24, 26^. Notably, OP9 stromal cells engineered to express DLL4 are able to support definitive hematopoiesis from arterial HE^24^. We attempted to obviate the need for feeder cells by including immobilised recombinant DLL4 during the EHT culture phase (Figure 1a). When we transferred the HSPC generated on DLL4 into conditions that support T cell development, we observed a striking improvement in progenitor T cell differentiation compared to HSPC generated on uncoated plates. The presence of DLL4 during EHT increased the median frequency of CD7+ cells generated after 7 days in downstream culture to 66.51% compared to 4.22% in the uncoated condition (P = 0.002, Mann-Whitney U test, Figure 1c). DLL4 increased the yield of CD7+ cells produced per input CD34+ cell over 90-fold compared to the uncoated control (Figure 1c). Encouragingly, DLL4 led to the emergence of a distinct population of the later stage CD5+, CD7+ T cell progenitors, a population that was nearly absent from the uncoated condition (Figure 1b). These data demonstrate that the addition of recombinant DLL4 during a chemically-defined EHT phase drastically improves the production of hematopoietic cells with T lineage potential from hPSCs.

Next, we attempted to improve progenitor T cell output even further. We previously demonstrated that the cell adhesion protein VCAM1 can increase the magnitude of notch signalling imparted by immobilised DLL4 during in vitro T cell differentiation^28^. We asked whether we could apply this principle to increase notch signalling during EHT and thus increase T cell potency.

When we supplied VCAM1 in combination with DLL4 during EHT, we observed a dramatic increase in downstream production of CD5+, CD7+ T cell progenitors (Figure 1b). VCAM1 and DLL4 together increased both the frequency (4.04 fold, P= 0.0022, Mann Whitney test) and yield per-input CD34+ cell (5.73 fold, P=0.0022, Mann Whitney test) of CD5+, CD7+ cells compared to DLL4 alone (Figure 1c). VCAM1 alone did not substantially alter progenitor T cell production compare to the uncoated control (Figure 1b,c), suggesting that it acts cooperatively with DLL4 to enhance the differentiation. This effect cannot be attributed to an increase in the number of non-adherent hematopoietic cells produced during EHT, as this number did not differ substantially between coating conditions (Supplementary Figure 1).

We tested whether T cell progenitor yields are improved by this EHT culture phase, compared to the method presented by Trotman-Grant et al, where CD34-enriched cells harvested from aggregates are placed directly into conditions that support T cell differentiation^16^ (Figure 1d). In agreement with previous reports, we found that >90% of CD34+ cells harvested from the day 8 aggregates generated in our protocol were CD43-, indicating that they have not yet established a hematopoietic identity^21^ (Figure 1e). In contrast, >95% of the non-adherent cells harvested after EHT co-expressed CD34 and CD43 (Figure 1e). Consistent with this transition to hematopoietic identity, the EHT phase improved the yield of CD5+, CD7+ T cell progenitors by 70-fold compared to seeding CD34+ cells directly into T cell differentiation conditions (Figure 1e,f). The EHT phase significantly improved T cell progenitor yield across multiple cell seeding densities (Figure 1f, P = 0.001, two-way ANOVA).

We conclude that an engineered signalling environment comprising recombinant DLL4 and VCAM1 and an appropriate chemically defined, serum-free media is sufficient to support emergence of HSPC with robust T lineage potential.

### DLL4 shifts hPSC-derived hematopoietic cell fate outputs

The addition of immobilized DLL4 and VCAM1 during EHT markedly impacted the ability of PSC-derived HSPC to differentiate towards the T lineage. We used droplet-based single cell RNA sequencing (scRNA-seq)^31^ to understand how these two proteins altered the HSPC transcriptional composition (Figure 2a). We compared non-adherent cells generated during EHT on uncoated plates, plates coated with DLL4 or VCAM1 alone, or coated with both proteins. Following doublet and dead-cell removal, we obtained 7,589 cells (Figure 2b) including at least 1,500 from each of the four coating conditions (Figure 2c). We performed unbiased clustering and differential gene expression analysis^32^ (Supplementary Figure 2). We detected a small population of stromal/mesenchymal cells that we excluded from downstream analysis (Supplementary Figure 2). We annotated the remaining four clusters as sub-types of HSPC on the basis of differential gene expression; *SPN*+ (CD43), *CD34*+ hematopoietic stem cells/multipotent progenitors (HSC/MPP), HLA class II+, *IFI16*+ myeloid progenitors, *HBD*+, *ITGA2B*+ erythroid/megakaryocyte progenitors and *SRGN*+, *MPO*+ neutrophil progenitors (Figure 2d, Supplementary Figure 2). Most cells expressed the hematopoietic gene *SPN* while the endothelial genes *KDR* and *CDH5* were nearly absent, consistent with successful transition from an endothelial to hematopoietic transcriptional program (Figure 2d, Supplementary Figure 2). Though *CD34* expression varies across cells at the RNA level (Supplementary Figure 2), the nearly ubiquitous cell surface expression of CD34 as measured by flow cytometry (Figure 1e) motivated us to label these hematopoietic sub-populations as progenitors.

**Figure 2:**
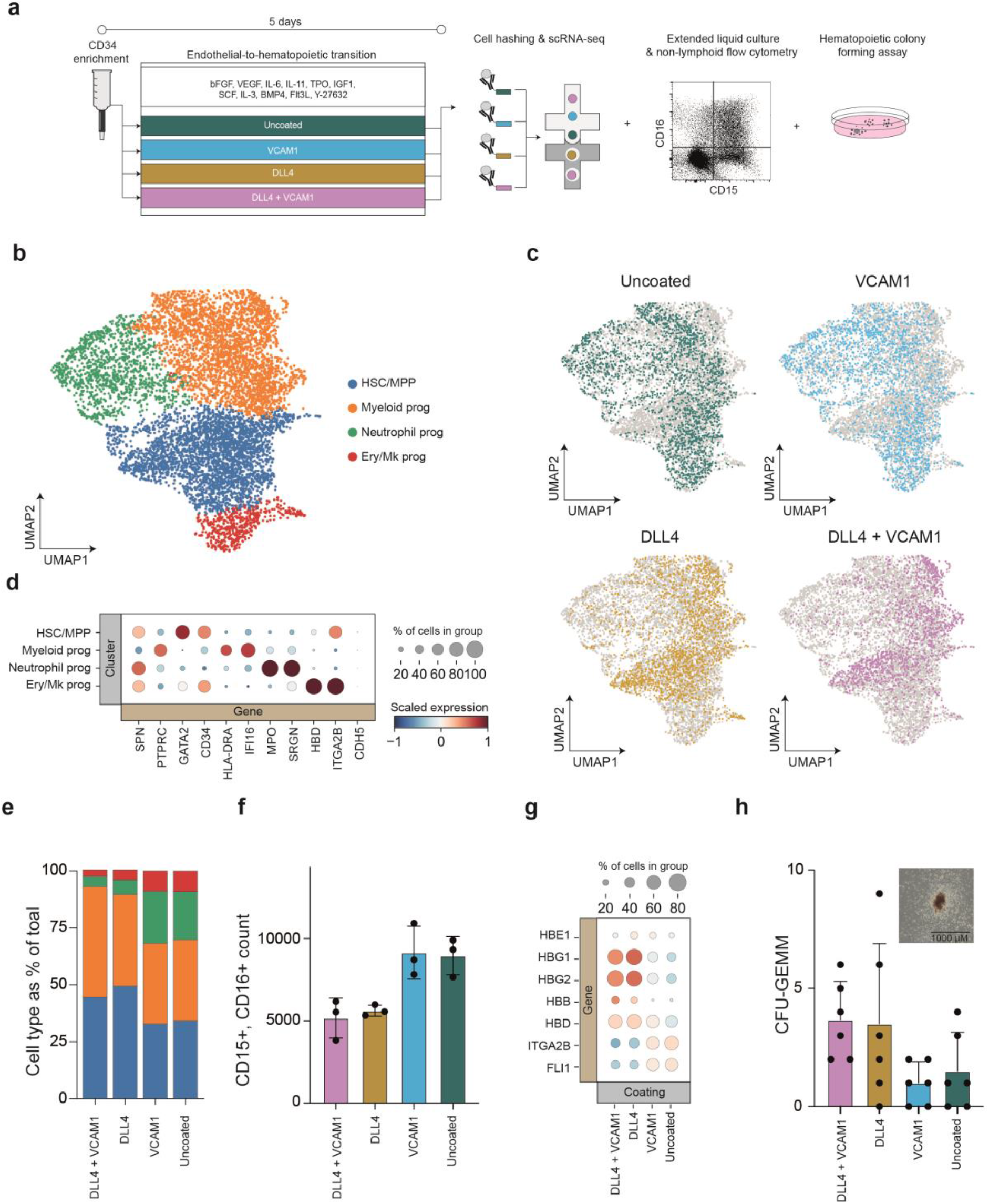
Engineered notch signalling during EHT reduces neutrophil differentiation and promotes definitive haematopoiesis. **a.)** Schematic overview of the experimental design used to test how the presence of DLL4 and VCAM1 during EHT impacts the resulting HSPC. **b.)** UMAP projection of cells identified in scRNA-sequencing after quality control filtration. Cells are coloured by unsupervised Leiden clusters annotated by expression of known marker genes. **c.)** UMAPs as in (b), coloured by coating condition. **d.)** Expression of hematopoietic and endothelial genes by cluster. Expression is scaled across all cells to a mean of 0 and unit variance. **e.)** Proportion of each hematopoietic progenitor subtype identified in scRNA-seq data, quantified for each coating condition. **f.)** Flow cytometry quantification of neutrophil output from HSPC generated in each EHT coating condition following extended 7-day liquid culture in myeloid and erythroid supportive media. **g.)** Comparison by coating condition of expression of haemoglobin genes and megakaryocyte-associated transcription factors within the MK/erythroid progenitor cluster. Expression is scaled across all cells to a mean of 0 and unit variance. **h.)** CFU-GEMM quantification from HSPC generated in each EHT coating condition following 14-day culture in semi-solid media (mean +/- s.d., n=6). Representative colony image shown is inset.

Coating conditions during EHT shifted the relative proportions of the resulting HSPC sub-types (Figure 2e). The presence of DLL4 caused an increase in the frequency of HSC/MPP, decreased the occurrence of erythroid/megakaryocyte progenitors and strongly reduced the abundance of neutrophil progenitors compared to VCAM1 alone or the uncoated control (Figure 2e). Upon extended liquid culture, we confirmed that DLL4 reduced the output of CD15+, CD16+ neutrophils 1.68-fold (Figure 2f, p = 0.0006, 2-way-ANOVA).

Within the erythroid/megakaryocyte progenitor population, we observed that DLL4 reduced expression of the megakaryocyte transcription factors I*TGA2B* and *FLI1*, and increased expression of haemoglobin genes associated with erythroid specification (Figure 2g). When we sub-clustered these cells, we observed an increase in the ratio of *KLF1*-expressing erythroid to *PLEK*-expressing megakaryocyte progenitors^33^ in the presence of DLL4 (Supplementary Figure 3). Extended liquid culture verified that DLL4 increased the ratio of CD235a+ erythroid cells to CD41+ megakaryocytes by 2.18-fold (Supplementary Figure 4, p<0.0001, 2-way-ANOVA).

Notch signalling during EHT *in vitro* has been reported to promote emergence of HSPC comparable to a later stage in human ontogeny as evidenced by a switch in globin gene expression from embryonic to foetal^24^. We examined globin gene expression within the erythroid/megakaryocyte cluster (Figure 2g). Indeed, upon addition of DLL4, we observed increased expression of the foetal globin genes *HBG1* (Log_2_ fold-change = 2.31, P_adjusted_ = 0.000031, Mann Whitney U Test) and *HBG2* (Log_2_ fold-change = 2.40, P_adjusted_ = 0.000069, Mann Whitney U Test) (Figure 2g).

Colony forming assays demonstrated that DLL4 increased the frequency of the multi-lineage granulocyte, erythrocyte, monocyte, megakaryocyte colony forming unit, CFU-GEMM by 2.9-fold (Figure 2h, p = 0.013, 2-way ANOVA), without significantly altering CFU-E and CFU-GM output (Supplementary Figure 4, p_CFU-E_ = 0.065, p_CFU-GM_ = 0.91, 2-way-ANOVA). DLL4 drives a shift in HSPC cell composition, including a reduction in neutrophil output, higher levels of foetal and adult to embryonic haemoglobin expression and an increase in the frequency of multipotent CFU-GEMM. While VCAM1 substantially enhanced T cell progenitor differentiation, this protein did not substantially alter the proportions of HSPC sub-types. To understand how the presence of DLL4 and VCAM1 during EHT influence lymphoid competency, we analysed transcriptional changes within the HSC/MPP cluster, the cell type from with lymphoid progenitors arise.

### DLL4 and VCAM1 alter HSC/MPP gene expression programs and cooperatively activate Notch signalling

We examined how the engineered signalling environment impacted the transcriptional identity of HSC/MPP produced during EHT. Expectedly, the presence of DLL4 increased the frequency and magnitude of expression of known downstream targets^34–37^ of Notch signalling (Figure 3a,b). In a separate developmental context, we previously showed that the cell adhesion molecule VCAM1 can enhance DLL4-mediated notch signalling^28^. VCAM1 alone did not substantially alter expression of notch targets (Figure 3b). In combination with DLL4, VCAM1 markedly increased expression of *HES1*, *CD3D*, *HES4*, *DTX1*, *BCL11B* and *HEY2* compared to DLL4 alone (Figure 3a, b, 1.35-fold increase in notch activity score, P = 3.36×10^-12^, Mann Whitney U Test). During EHT, VCAM synergises with DLL4 to promote high levels of notch activity in HSC/MPP (2-way ANOVA, interaction p < 0.0001).

**Figure 3:**
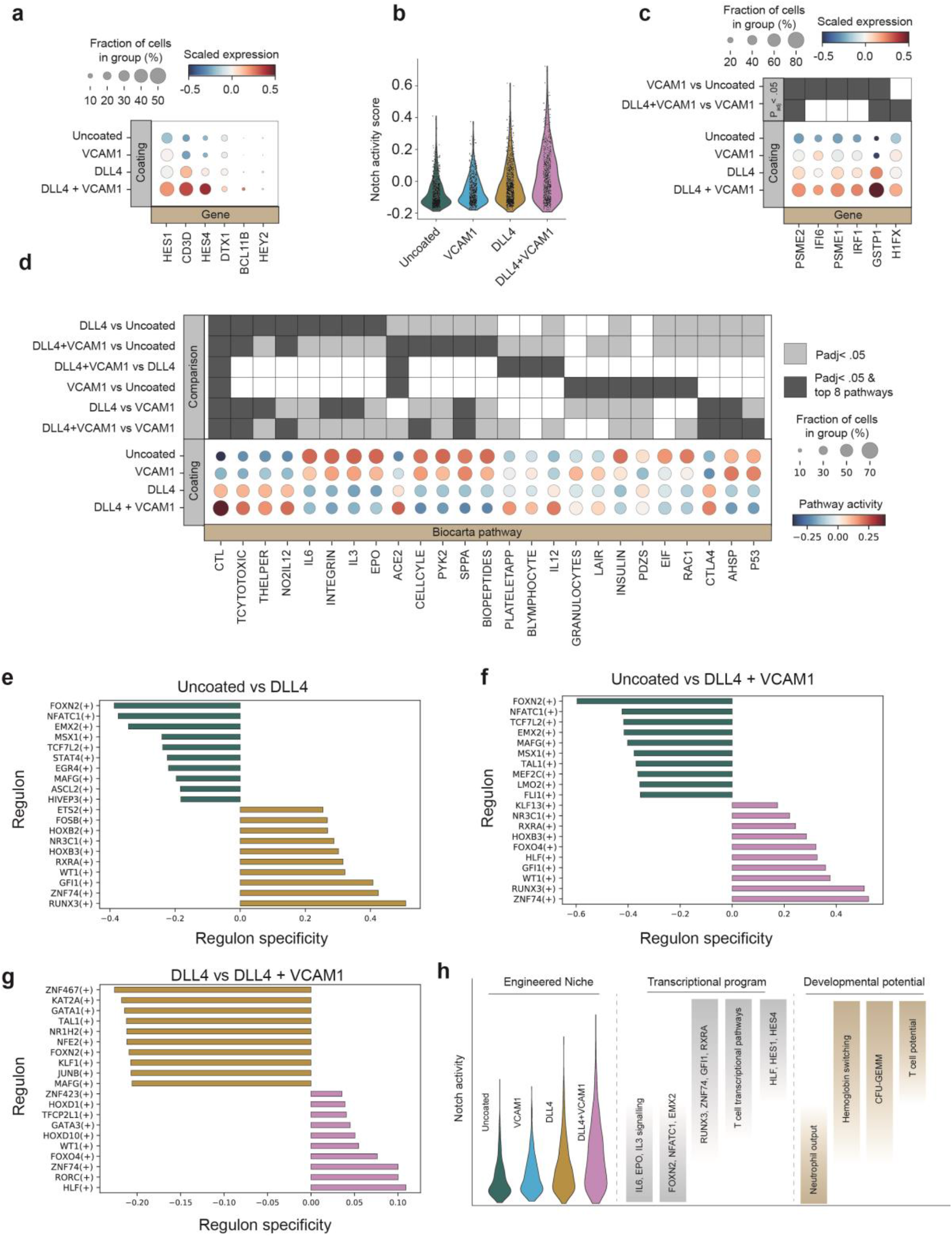
VCAM1 promotes an inflammatory program and cooperates with DLL4 to enhance notch signalling and hematopoietic gene expression in HSC/MPP. **a.)** Expression of known notch target genes were analysed in cells within the HSC/MPP cluster from each coating condition. Expression of each gene scaled to a mean of 0 and unit variance within all HSC/MPP. **b.)** The geometric means of the notch target genes shown in (a) were combined on a per-cell basis to create a single-cell notch activity score. **c.)** Expression of genes that are impacted by VCAM1 within the HSC/MPP cluster. Differential expression analysis (Wilcoxon rank sum) was performed between VCAM1 and uncoated as well as between VCAM1+DLL4 and DLL4 alone. The 20 genes with the highest z-score from each comparison were retained and filtered for P_adjusted_ <0.05. All genes meeting these criteria are plotted. **d.)** Differential pathway activity within HSC/MPP across coating conditions. Pairwise comparisons were performed between each coating condition and all others and the top 8 differentially active pathways by z-score from each comparison are depicted. Box shading indicates which pathways are in the top 8 of a given comparison and which comparisons have P_adjusted_ <0.05. **e, f, g.)** Comparison of SCENIC regulon activity between coating conditions. The top 10 regulons specific to each coating condition are shown. Regulon specificity is calculated as the Jensen-Shannon distance between conditions. **h.)** Model of the impact of coating conditions on notch signalling and associated notch dependence of transcriptional programs and cell fate decisions.

Next, we undertook an unbiased exploration of the impact of coating conditions on MPP/HSC gene expression (Figure 3c, d, Supplementary Figure 5). In addition to increasing Notch signalling, differential expression analyses revealed that VCAM1 also altered expression of *EVL* and *FERMT3*, genes involved in cell adhesion and cytoskeletal polymerisation^38, 39^ (Supplementary Figure 5), and increased activity of the interferon induced genes *IFI6* and *IRF1* as well as members of interferon-induced immunoproteosome, *PSME2* and *PSME1* (Figure 3c). Changes in pro-inflammatory, interferon responsive genes were observed, both when comparing VCAM1 to the uncoated control, and when comparing VCAM1 and DLL4 to DLL4 alone (Figure 3c).

Pathways associated with lymphoid and T cell identity were amongst the most strongly activated by DLL4 (Figure 3d). CTL, TCYTOTOXIC, THELPER were all highly upregulated by DLL4 and these effects were magnified by the addition of VCAM1 (Figure 3d). DLL4 reduced activity of cell cycle and P53 pathways and increased activity of the T cell inhibitory pathway CTLA4. These alterations in transcriptional state are consistent with a model whereby notch signalling promotes emergence of an HSC/MPP population that is primed to undergo TCR-mediated selection during T cell differentiation.

SCENIC analysis^40^ revealed regulons (transcription factors and their downstream targets) that were specifically upregulated in each engineered signalling environment (Figure 3e,f,g). Nearly all of the regulons that were most strongly activated by DLL4 have a known role in blood emergence and T cell differentiation in human or mouse (Figure 3e). *RUNX3* contributes to HSC maintenance and T cell differentiation^41, 42^. *GFI1* and *FOSB* are important drivers of endothelial to hematopoietic transition^43, 44^. The homeobox gene *HOXB3* is expressed in both uncommitted HSCs and during T cell development^45^. *WT1* is important for survival and maintenance HSC/MPP and downregulated in differentiated progeny^46, 47^. The DLL4-mediated increase in *RXRA* regulon activity merits future investigation given the complex role of retinoic acid signalling during the emergence of HSCs from hPSCs^48–50^. *ZNF74* was also amongst the regulons most strongly upregulated by DLL4 and has no previously documented role in hematopoiesis (Figure 3e,f).

Notably, when added alongside DLL4, VCAM1 increased *HLF* and *GATA3* regulon activity (Figure 3g). *HLF* expression is highly correlated with definitive HSC identity in human cord-blood, foetal liver, peripheral blood and bone-marrow^51^. In addition to its appreciated role in human T cell development, *Gata3* helps maintain HSC identity in mice^52^.

Collectively, these data demonstrate that introducing DLL4 and VCAM1 during EHT positively shifts the distribution of notch pathway activity in the resulting HSC/MPP (Figure 3b, h). This shift in notch activity is associated with marked changes in transcription factor and pathway activity and alters cell fate distribution, with intermediate levels of notch activity driving haemoglobin switching and suppressing neutrophil output and high notch activity unlocking T cell potential (Figure 3h).

### Hematopoietic progenitors generated in the presence of DLL4 and VCAM1 are capable of maturation into CD8+ T cells

After demonstrating that the addition of DLL4 and VCAM1 during EHT is sufficient to promote emergence of CD7+, CD5+ T cell progenitors, we sought to verify that these cells are able to develop into CD4+, CD8+ DP cells and CD8+, CD4-, CD3+, TCRαβ+ mature T cells. (Figure 4a-g). We transferred PSC-derived hematopoietic progenitors into a serum-free media supplemented with a cytokine composition that we previously optimised for generating CD4+, CD8+ DP cells from umbilical cord blood (UCB)^53^ (Figure 4b). Unfortunately, this media did not efficiently support T cell maturation from hPSCs. (Figure 4f,g). We thus set out to understand how the PSC-derived HSPC produced in our protocol respond to different cytokine concentrations over time, and use that information to develop an optimised PSC-derived T cell maturation method.

**Figure 4.**
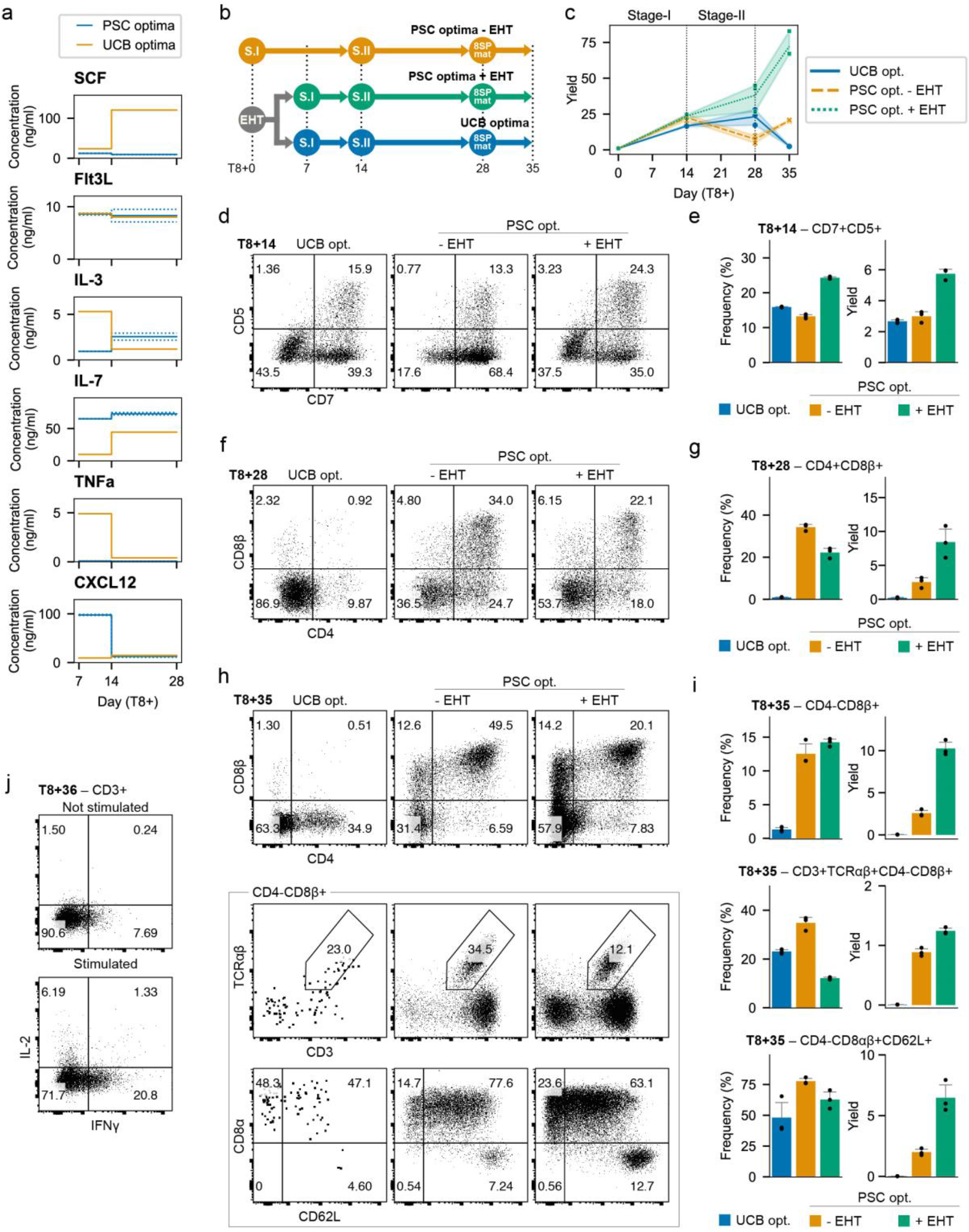
Optimized cytokines enhance T-cell development. **a.)** Predicted optima for PSC-derived T-cell differentiation compared with previously identified optima for umbilical cord blood (UCB)-derived HSPC differentiation to the T-lineage. **b.)** To validate predictions, CD34+ cells were cultured in EHT conditions for 7 days then transferred to cytokines optimized for PSC- or CB-derived cells. Additionally, CD34+ were placed directly into PSC optima without the EHT step. **c.)** Total viable cell yield for each test condition per input CD34+ cell (n = 3). **d.)** Representative flow cytometry of CD7+,CD5+ T cell progenitors at day 8 + 14. **e.)** Quantification of the frequency and yield of CD7+,CD5+ T cell progenitors (yield is reported as number of CD5+,CD7+ cells per CD34+ input cell, n=3, mean +/- s.d.). **f.)** Representative flow cytometry of CD4+,CD8+ DP cells at day 8 + 28. **g.)** Quantification of the frequency and yield of CD4+,CD8+ DP cells (yield is reported as number of CD4+,CD8+ cells per CD34+ input cell, n=3, mean +/- s.d.). **h.)** Representative flow cytometry at day 8+35 after stimulation with anti-CD2/3/28 antibodies in the presence of + IL-21. **i)** Quantification of the frequency and yield of the phenotypes shown in (h). Yield is reported as abundance of indicate cell population per CD34+ input cell (n=3, mean +/- s.d). **j.)** Flow cytometry of differentiation cultures after stimulation with PMA and ionomycin. Representative of 3 replicates.

We have previously shown that statistical dose response models are highly effective for optimizing stage-specific differentiation media formulations^53^. We conducted a 6 cytokine, 5 concentration factorial-type experiment over two stages of T cell development (Supplementary Figure 6). In the first 7 days post-EHT, we measured CD5+,CD7+ proT, CD4+,CD8-,CD3-immature single positive (CD4ISP), and CD4+,CD8+,CD3-DP. In the subsequent 14 days we measured CD3-DP, CD3+ DP, and CD4-, CD8+, CD3+ (CD8SP). An equal number of cells were seeded in each test condition and we measured the absolute number of each cell-type at the end of each stage using flow cytometry. We then used this information to derive polynomial dose response models using regression.

Over the 7-day early differentiation stage, proT, CD4ISP and DP, CD3+ output had a strong positive response to IL-7 concentration and a moderate positive response to CXCL12. Outputs of these populations exhibited a negative response to increasing TNFα concentrations. During the maturation stage, CD3- and CD3+DP as well as CD8SP responded positively to IL-7 and IL-3 and negatively to TNFα (Supplementary Figure 6). This experimental design also allowed us to examine multiplicative interactions between cytokines (Supplementary Figure 6).

Next, we sought to determine an optimal combination of cytokine concentrations for each differentiation stage. We defined a desirability function to maximize the number of ProT, CD4ISP and CD3-,DP cells for the early stage and CD3-,DP, CD3+,DP and CD8SP for the maturation stage (Supplementary Figure 6). Each desirability function scales the predicted output from our dose response models between [0,1] for a given set of cytokine concentrations. We then combined the value of each desirability function using the geometric mean to calculate an overall desirability score that could be optimized using single objective optimization algorithms. Each stage was optimized separately using the basin-hopping algorithm^54^. To avoid settling in local optima, we repeated the procedure with 25 random initializations and retained the top five most desirable solutions. The cytokine concentrations that elicited these five responses were then averaged to provide a set of optimal cytokine concentrations for each stage of T cell development from PSC-derived hematopoietic progenitors (Supplementary Figure 6, Figure 4a).

We compared these two newly optimised media formulations (PSC optima) against the previously developed cord blood-optimised media^53^ (UCB opt, Figure 4a,b). PSC optima improved total cellularity compared to the CB control as early as day 14 and the magnitude of this effect was amplified over the course of the differentiation (Figure 4c). Furthermore, the yield of desired cell types was drastically improved by the new media formulation. The PSC-optimised early stage media increased the abundance of CD7+, CD5+ proT cells two-fold compared to the CB control (Figure 4d,e).

Following the maturation stage, PSC optima caused a striking 40-fold improvement in CD4+, CD8+ DP yields compared to the CB control (Figure 4f,g). We non-specifically activated TCR signalling with antibodies against CD3, CD28 and CD2 to simulate positive selection and quantified the yield of CD8SP, CD3+, TCRαβ+cells. The optimised media improved the yield by more than 150-fold compared to the UCB optima (Figure 4h,i).

One limitation of previous *in vitro* differentiation methods, is that they tend to produce T cells with unconventional immunophenotypes. A common unintended product are cells that express a CD8αα homo-dimer, a characteristic feature of innate-like cells^10, 14^. PSC-derived T cells have previously been reported to lack robust expression of the adhesion molecule CD62L^10, 14^. The T cells generated in our protocol express the conventional CD8αβ heterodimer and robustly express CD62L (Figure 4h,i). They produce IFNγ and IL-2 in response to non-specific stimulation using PMA/ionomycin (Figure 4j). Collectively, these data demonstrate that PSC-derived hematopoietic progenitors generated in the presence of DLL4 and VCAM1 and capable of efficient differentiation, not only into T cell progenitors, but into mature and functional CD8SP, CD3+, TCRαβ+cells.

The engineered EHT niche and optimised T cell differentiation media that we establish here each substantially improve T cell production. Collectively, these improvements contribute to a highly efficient protocol that will aid both developmental studies and scalable T cell manufacturing for cell therapy applications.

## Discussion

In this study, we present a substantially improved serum- and feeder-free method for generating hematopoietic progenitors and T cells from hPSCs. Two other feeder-free hPSC-to-T cell differentiation systems have been reported to date^13, 16^. The method presented by Iriguchi et al. does not support endogenous αβTCR recombination, which is an important limitation that precludes research into key stages of T cell development^13^. The protocol presented by Trotman-Grant et al. supports αβTCR recombination, but lacks an explicit EHT culture phase^16^. Here we find that the addition of an EHT stage comprising DLL4 and VCAM1 drastically improves progenitor T cell differentiation. Furthermore, the addition of the EHT stage faithfully recapitulates the distinct steps and physical niches required for HSPC emergence and T cell differentiation as they occur in human ontogeny^22, 55^.

Here we verify previous reports that notch signalling during EHT is critical for emergence of HSC/MPP with T-lineage potential^24, 26^. While DLL4 might also impact HSC/MPP once they have already emerged, notch inhibition experiments demonstrate that this pathway is indispensable during the EHT process itself^26^. We do not see any evidence of T cell progenitors emerging during our EHT phase which lacks the necessary cytokine IL-7^56^. Rather, we observe a definitive HSC/MPP population that is capable of lymphoid differentiation.

We show that VCAM1 cooperates with DLL4 to enhance the magnitude of notch signalling during the EHT culture phase. Unexpectedly, we also found that VCAM1 promotes inflammatory-responsive gene expression in nascent HSC/MPP. Studies in zebrafish and mouse have shown that inflammatory signals are important for HSC emergence, both upstream and downstream of the notch pathway^57–60^. The differentiation method we present here will enable future interrogation into the role of inflammatory signalling during human EHT.

When we first attempted to differentiate hPSC derived T cell progenitors into mature T cells using a media that we previously optimised for cord-blood derived progenitors, we were largely unsuccessful. Upon re-optimising the media for hPSC derived cells, we observed >150-fold increase in mature T cell production. Notably, the PSC optimised media uses the exact same cytokines as the cord-blood media and these formulations differ only in their concentrations. These distinct optima may reflect differences in the developmental stage and transcriptional state of the starting material. PSC-derived cells required higher concentrations of CXCL12 which may be necessary to produce and maintain early lymphoid progenitors which are already abundant in cord-blood^61^. PSC-derived cells also required markedly less of the inflammatory cytokine TNFα compared to cord-blood. The inflammatory program promoted by VCAM1 during EHT may lessen the subsequent requirement for inflammatory cytokines in the media.

Interestingly, the early T cell differentiation media that we optimised for PSCs can partially replace the EHT media, albeit at drastically reduced efficiency. Future work will determine whether the early T cell media can actually promote EHT of HE cells, or whether it simply allows the very small number of hematopoietic cells present in the day 8 aggregates to efficiently differentiate towards the T lineage.

The method we report here has widespread applicability. We have shown that a defined EHT process allows us to study the impact of juxtracrine factors on the transcriptome and lineage potential of emergent hematopoietic stem and progenitor cells. In another application, we used this method to study how cytokine dosages impacts T cell maturation. Though we developed this platform to achieve robust, efficient and customisable T cell differentiation from hPSC, we have shown that this method can be used to produce and study multiple hematopoietic cell types. In addition to producing T cell progenitors, the defined EHT process also generates erythroid/megakaryocyte and myeloid progenitors and will enable basic research into blood stem cell emergence and lineage specification.

## Methods

### Human Pluripotent Stem Cell (hPSC) Culture

The human induced pluripotent stem cell line iPS11(Alstem Cell Advancements, episomally derived from human foreskin fibroblasts) was cultured on tissue culture-treated plasticware pre-coated with 1mL per well Geltrex (Life Technologies A1413302) for 1 hour at 37C. Cells were cultured in serum-free media (mTeSR1, Stemcell technologies 85850) that was supplemented with Penicillin-Streptomycin (Invitrogen, 15140122-0.5% V/V). Growth media was aspirated and replenished daily and cells were maintained at 37C, 5% CO_2_. Following thawing or passaging, media was additionally supplemented with the ROCK inhibitor Y-27632 (Stemcell Technologies, 72308) at 7.5uM and 5uM concentrations respectively. To passage, cells were partially dissociated to small aggregates with 1x TrypLE Express (Thermofisher 12605028) at 37 C followed by cell scraping. Dissociation was carried out for 2 to 4 minutes.

### Aggregation and CD34+ induction

For CD34+ induction media formulations, see Supplementary Workbook 1. iPS11 cells were grown to ∼90% confluency and dissociated to single cells with TrypLE Express for 3 to 5 minutes at 37 C. Dissociated cells were collected and counted and a desired cell number was pelleted at 200xg for 5 minutes. Following supernatant aspiration, cells were resuspended in 2mL per well T0 media and deposited into AggreWell 400 6-Well plates (StemCell Technologies 34425) prepared with Aggrewell Rinsing Solution (StemCell Technologies 07010) according to the manufacturer’s instructions. Cells were seeded at a density of 180 cells per microwell and aggregated by centrifugation at 200xg for 5 minutes. For the duration of the CD34+ induction, cells were cultured at 37 C in a hypoxia incubator at 5% CO_2_, 5% O_2_.

24 hours after seeding, 2mL of T1 media was added to each well. 42 hours after initiating the differentiation, media was aspirated and replaced with 2mL per well T1.75 media. The following day, an additional 2mL of T1.75 media was added to each well. 96 hours after initiating the differentiation, the T1.75 media was aspirated and replaced with 2mL/well T4 media. 48 hours later, the media in each well was supplemented with 2mL of T6 media. The following day, a subset of aggregates was harvested and dissociated with TrypLE Express, washed with 200uL HF buffer (HBSS, Thermo Fisher Scientific, 14175103, supplemented with 2% Fetal bovine serum, Thermo Fisher Scientific, 12483020) and stained with antibodies against CD34, CD43, CD73 and CD184 and analysed by flow cytometry.

192 hours after initiating the differentiations, aggregates were collected and pelleted by centrifugation at 200xg for 5 minutes. Spent media was aspirated and aggregates were dissociated in 3mL TryPLE supplemented with DNaseI (Millipore Sigma, 260913-10MU) for 10 to 15 minutes. Cells were pipetted to a single cell suspension, washed and counted. CD34+ cells were enriched using the CD34 positive selection kit (Miltenyi Biotec, 130-046-702) according to the manufacturer’s instructions. CD34+ enriched cells were cryopreserved using CryoStor CS10 (StemCell Technologies, 07930) for use in downstream culture.

### Endothelial-to-hematopoietic transition (EHT) culture

CD34+ cells generated above were used as input for EHT culture. Coating solution was prepared using sterile PBS combined with 15ug/mL Fc-tagged DLL4 (Sino Biological, 10171-H02H) and 2.5ug/mL Fc-tagged VCAM1 (R&D Systems, 643-VM). Tissue culture-treated 96 well plates (Fisher Scientific, 12-556-008) were pre-coated with 50uL of coating solution overnight at 4 C. Coating solution was aspirated and plates were washed with PBS immediately prior to use. CD34+ enriched cells were resuspended in EHT media (Supplementary Workbook 1) at a concentration of 1×10^5^ cells per mL unless otherwise indicated. 100uL (10,000 cells) were seeded on to each well of the 96 well plate unless otherwise indicated. Cultures were incubated at 37C, 5% CO_2_ for 5 or 7 days as specified in the main text and non-adherent cells were harvested by gentle pipetting.

### T cell differentiation

Non-adherent cells harvested from EHT cultures were pelleted by centrifugation at 300xg for 5 minutes. For the data shown in Figure 1, spent media was aspirated and cells were re-suspended in an un-optimised T cell differentiation media at a split at a ratio of 1:3. Un-optimised T cell differentiation media was prepared by combining IMDM with GlutaMAX basal media (Thermo Fisher Scientific, 31980030) with 20% BIT 9500 serum substitute (STEMCELL Technologies, 09500) 0.05% LDL(STEMCELL Technologies, 02698), 60 µM Ascorbic acid (Sigma-Aldrich, A8960), 24µM 2-Mercaptoethanol (Sigma-Aldrich, M3148), 0.02µg/mL SCF (R&D Systems, 255-SC), 0.02µg/mL Flt3L (R&D Systems, 308-FK), 0.02µg/mL TPO (R&D Systems, 288-TP), 0.02µg/mL IL-7(R&D Systems, 207-IL), 0.01µg/mL IL-3 (R&D Systems, 203-IL) and 0.005µg/mL TNFα (R&D Systems, 210-TA). Cells were seeded in 100uL per well of a 96 well plate pre-coated with DLL4 and VCAM1 as described above. 3 to 4 days after seeding, cells were fed with an additional 100uL per well of un-optimized T cell differentiation media. 7 days after seeding, cells were harvested by pipetting and seeded on to freshly coated plated in un-optimized T cell differentiation media and cultured for an additional 7 days, with a media top-up after 3 days.

For the experiments described in Figure 4, cells were cultured as described above but using either cord-blood optimised early and late T cell differentiation media or PSC-optimised early and late T cell differentiation media (Supplementary Workbook 1). After EHT, cells were split 1:3 and seeded into 96 well plates pre-coated with DLL4 and VCAM1 in 100µL per-well early T cell differentiation media for 7 days with a 100µL top-up on day 3 or 4. After 7 days, cells were split 1:3 into 96 well plates freshly coated with DLL4 and VCAM1 as described above in late T cell differentiation media where they were cultured for 14 days with a 100µL top-up on day 3 or 4 followed by half-media changes twice per week. All T cell differentiation cultures were carried at 37C, 5% CO_2_.

After 7 days of EHT culture and 21 days of T cell differentiation, T cells were stimulated to promote DP to CD8SP maturation. Stimulation was carried out in 96 well plates pre-coated with DLL4 and VCAM1 as described above in 100µL per well late T cell differentiation media supplemented with 20ng/mL IL-21 (R&D Systems, 8879-IL) and 1.25% ImmunoCult Human CD3/CD28/CD2 T cell activator (STEMCELL Technologies, 10970). After 48 hours, each well was topped up with an additional 100 µL late T cell differentiation media with IL-21 but without the T cell activator reagent where they were cultured for an additional 5 days.

### Extended myeloid and erythroid differentiation

Non-adherent cells harvested from day 5 EHT cultures were pelleted by centrifugation at 300xg for 5 minutes. Cells were counted using a hemocytometer and seeded at 2500 cells/well of a 96 well plate in 200 µL erythroid/myeloid supporting media (Iscove’s Modified Dulbecco’s Medium supplemented with Glutamax, ThermoFisher Scientific 31980030), 30% FBS, 1% bovine serum albumin (Wisent Bioproducts 809-098-EL), 0.1 mM 2-mercaptoethanol (Sigma-Aldrich M3148), 20 ng/mL IL-6 (R&D Systems 206-IL), 50 ng/mL SCF (R&D Systems 255-SC), 20ng/mL IL-3 (R&D Systems 203-IL), 20 ng/mL G-CSF (R&D Systems 214-CS), 20 ng/mL GM-CSF (R&D Systems 215-GM), and 3 U/mL EPO (R&D Systems 287-TC). Cells were cultured for 7 days followed EHT and assessed using by flow cytometry.

### Colony forming assays

Non-adherent cells harvested from Day 5 EHT cultures were pelleted by centrifugation at 300xg for 5 minutes. Cells were counted using a hemocytometer and seeded in 35 mm dishes at 500 cells/dish in MethoCult H4435 Enriched (StemCell Technologies 04435) according to the manufacturer’s instructions. CFU-GM, CFU-E and CFU-GEMM Colony identification was performed 14 days after seeding in MethoCult media according to the Atlas of Human Hematopoietic Colonies (StemCell Technologies, 28700).

### Flow cytometry

Samples were collected in 96 well V bottom plates and pelleted at 300 x g for 5 minutes. Supernatant was aspirated and cells were washed in 200 uL per well PBS (ThermoFisher 10010049). Cells were incubated for 15 minutes at room temperature in the dark in 50uL PBS with Fc Block (1/100 dilution, BD, 564220) and Zombie UV dye (1 in 500 dilution, BioLegend 423108). Next, 50 uL of antibody cocktail made up at 2x working concentration in HF buffer (HBSS, Thermo Fisher Scientific, 14175103, supplemented with 2% Fetal bovine serum, Thermo Fisher Scientific, 12483020) was mixed 1:1 with the cell suspension. Cells were stained for 30 minutes on ice protected from light and washed twice with 200uL per well HF buffer. Samples were analysed on a CytoFLEX LX cytometer (Beckman), compensation was performed in CytExpert v2.3 and data were gated and plotted in FlowJo v9. Antibodies used are listed in Supplementary Workbook 1.

### Sample preparation for scRNA-sequencing

EHT cultures were initiated and carried out as described above. Three wells of a 96 well flat bottom TC-treated plate were prepared for each coating condition and cells were cultured in EHT media for 5 days. On day 5, non-adherent cells were harvested by gentle pipetting. Well replicates were pooled for each condition and pelleted by centrifugation at 300xg for 5 minutes. Samples were washed with 200 uL HF buffer (HBSS, Thermo Fisher Scientific, 14175103, supplemented with 2% Fetal bovine serum, Thermo Fisher Scientific, 12483020). Cells were stained with 0.1ug TotalSeq-B anti-human Hashtag antibody (Biolegend, 394633, 394635, 394637 or 394639) per sample diluted in 100uL HF buffer for 30 minutes on ice. Cells were washed 3 times with HF buffer. Cell density was determined by hand counting using a hemocytometer and samples were pooled at equal cell ratios per sample. The pooled sample was counted and loaded on the chromium controller as per the manufacturer’s instructions aiming for a target cell capture of 5,900 cells.

### scRNA-sequencing analysis

The Cell Ranger^31^ function mkfastq was used for FASTQ generation, and Cell Ranger count was applied using default parameters for trimming, alignment and counting using hg19 as the reference genome. The data were further processed in python using the scanpy package ^32^ (Version 1.7.0). Demultiplexed sample antibody counts from the Cell Ranger pipeline were used to identify and filter out empty droplets and doublets. Doublets were then further identified and filtered by removing cells that had greater than 36000 counts or that had greater than 6000 genes. Dying/dead cells were removed by filtering cells that had a high fraction of mitochondrial reads (>18%). Genes that were expressed in few cells (n=3) were also removed from the processed dataset. The fully filtered dataset contained 7589 cells and 20546 genes. The expression matrix was then normalized to 1×10^4^ counts per cell and was log transformed. Highly variable genes were computed before performing principal component analysis (50 principal components), generating a neighbours graph, and performing leiden clustering to cluster cells. A UMAP embedding was generated for dataset visualization.

Cell cycle signatures in the dataset were annotated using a previously reported gene list^62^ and the G2_M and G1_S signatures were regressed using the scanpy regress_out function to remove cell cycle contributions to the expression matrix. Leiden clusters (resolution=0.18) were annotated using marker genes as indicated in the main text. Differential expression was performed with the scanpy package using a wilcoxon rank sum test^32^. Differential expression was performed between cell types, and between plating conditions within the multipotent progenitor cells. Reported P_adjusted_ values were calculated using Benjamini-Hochberg correction.

Regulons were computed with the pySCENIC pipeline^40^. Grnboost2 was used to determine coexpression modules and hg19 transcription factor motifs provided by the Aerts lab were used to identify direct transcription factor motif interactions. Regulon activity was quantified in single cells using the Area Under Curve metric built into the pipeline. Differential regulon activity was computed using the SCENIC regulon specificity score and with scanpy’s rank_genes_groups function ^32, 63^.

Pathway analysis was performed using the gseapy package’s (v0.10.5) single-sample geneset enrichment analysis (ssgsea) function and the Biocarta pathway database provided by the gseapy package^64, 65^. Reported P_adjusted_ values were calculated using Benjamini-Hochberg correction.

### Cytokine dose response models during T cell development

Cytokine dose responses were measured using a 6-factor orthogonal central composite design (CCD) experiment using JMP14 software. The CCD combines a two-level fractional factorial, where concentrations are scaled to (−1,1), with added centre (0) and axial (-α,α) points. The value of α=2.366 was chosen automatically by JMP14 to ensure orthogonality. This resulted in a five-level design capable of estimating all linear, quadratic, and two-factor interaction terms. Select third-order terms were also included where they improved model fit. Cytokine concentrations were scaled logarithmically to provide a continuous mapping between actual and scaled values. Experiments were laid out using an epMotion 5073 liquid handler (Eppendorf). For first stage, cells were passaged after EHT to test conditions at equal densities and cultured for 7 days. For the second stage, cells post-EHT were cultured with cytokine concentrations able to support T-lineage induction and cultured for 7 days. Subsequently, these cells were pooled and passaged to test conditions and cultured for an additional 14 days. At the end of each experiment, phenotype and absolute cell numbers were measured using flow cytometry. The number of cells in each population of interest were transformed as 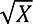 or log(*X* + 1) and dose response models were fit using least-squares regression. The concentrations of cytokines tested are provided in Supplementary Table 1. Calculated regression coefficients are provided in Supplementary Tables 2-7.

### Optimizing dose response models for generating mature T cells

Regression coefficient estimates were used to construct polynomial models using custom code written in Python 3.7. Desirability functions were applied to each model to maximize the response of that cell population and an overall desirability was calculated at the geometric mean of each individual desirability score^66^. The overall desirability was then optimized using the basin hopping algorithm^54^ from the *SciPy* Python library. The solution space was constrained to be within a hypersphere with radius α=2.366 to ensure that optimal solutions were not extrapolations beyond the range of cytokine concentrations tested. Within this space, the optimizer was randomly initialized 25 times and the top five most desirable solutions were averaged to provide an optimal set of cytokine concentrations (Supplementary Workbook 1). The custom code used for optimization is available at gitlab.com/stemcellbioengineering/polynomialfeatures (v1.1).

## Supporting information

Supplementary Workbook 1

## Acknowledgements

The authors would like to acknowledge Tara Stach, Yvonne Chung and Bernie Zhao from BRC-seq for their technical support. We thank Ashton Trotman-Grant and Juan-Carlos Zúñiga-Pflücker for valuable guidance. Y.S.M is supported by a Michael Smith Foundation for Health Research Trainee award and the Canadian Institute for Health Research (CIHR) Banting Fellowship. M.C.M was funded by a Natural Sciences and Engineering Research Council of Canada (NSERC) USRA and CBR summer award. J.M.E is funded by a NSERC Canadian Graduate Scholarship and Zymeworks-Michael Smith Laboratories Fellowship in Advanced Protein Engineering. E.L.C holds a CIHR Vanier scholarship. P.W.Z is a Canada Research Chair in Stem Cell Bioengineering. We acknowledge funding from the Stem Cell Network Fuelling Biotechnology Partnerships award, Notch Therapeutics, a CIHR Foundation Grant FRN (154283) and from the Wellcome Leap Human Organs, Physiology, and Engineering (HOPE) program.

## Contributions

Y.S.M. and P.W.Z designed the study. Y.S.M., J.M.E., E.L.C., C.Z., T.Y., A.H. & C.L., conducted the experiments. Y.S.M., J.M.E., M.C.M., I.I.R. & D.J.H.F.K analysed the data. Y.S.M, J.M.E. & P.Z. prepared the manuscript. All authors reviewed the manuscript.

## Ethics declaration

Y.S.M., P.W.Z and J.M.E are inventors of patents relating to T cell differentiation from stem cells. A.H. is currently employed by Notch Therapeutics, a T cell therapy company. Y.S.M and J.M.E. consult for regenerative medicine companies. Funding for this work was provided by Notch Therapeutics.

