## Supplementary Information for "DLL4 and VCAM1 enhance the emergence of T cell-competent hematopoietic progenitors from human pluripotent stem cells"

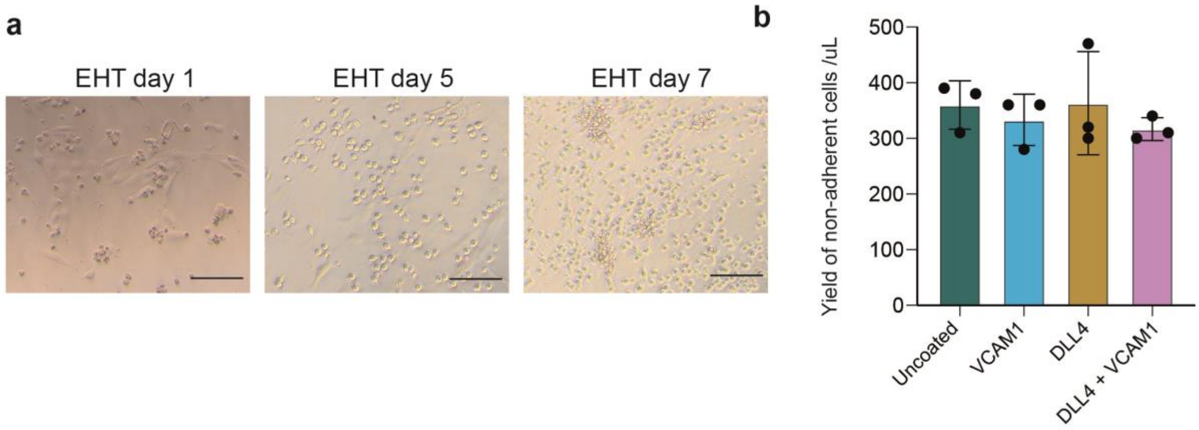


**Supplementary Figure 1: Endothelial to hematopoietic transition of PSC derived CD34+ cells. a.)** Bright field images of PSC derived CD34+ cells seeded into EHT cultures in the presence of immobilised DLL4 and VCAM1at the timepoints indicated. Scale bars = 100uM) **b.)** The yield of non-adherent hematopoietic cells was quantified after 5 days in EHT under the coating conditions indicated.

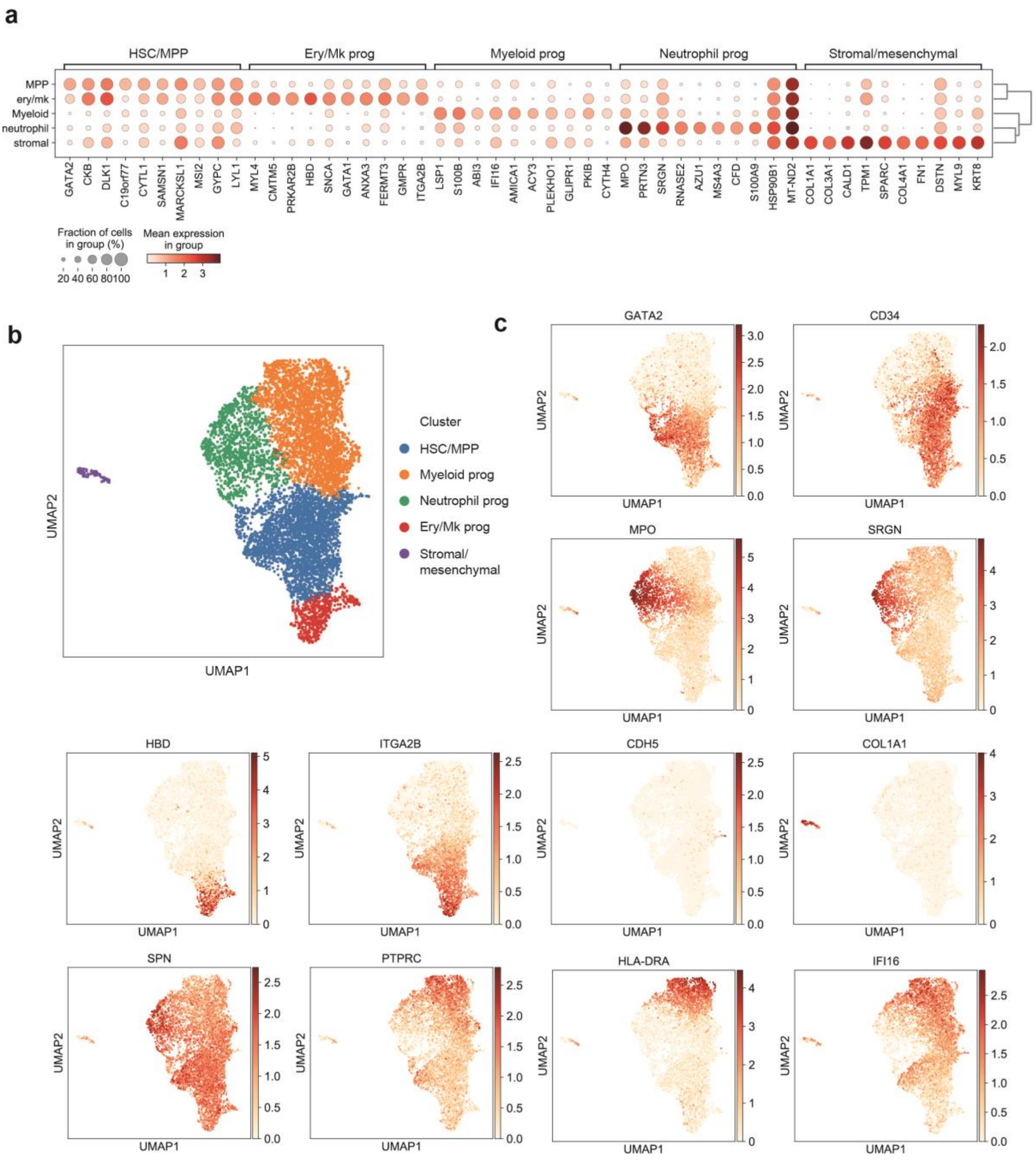


**Supplementary Figure 2: Identification of cell types generated during EHT from PSC-derived CD34+ cells. a.)** Non-adherent cells that emerged during EHT were collected and analysed by scRNA-sequencing. We performed unsupervised Leiden clustering and identified 5 cell clusters. Next, we identified genes that were differentially expressed between cells of each cluster and all other cells (Mann Whitney U-test). The 10 genes with the highest Z score for each cluster are plotted. **b.)** UMAP projection colored by cluster annotation. **c.)** UMAP projection for a selection of marker genes characteristic of each cluster.

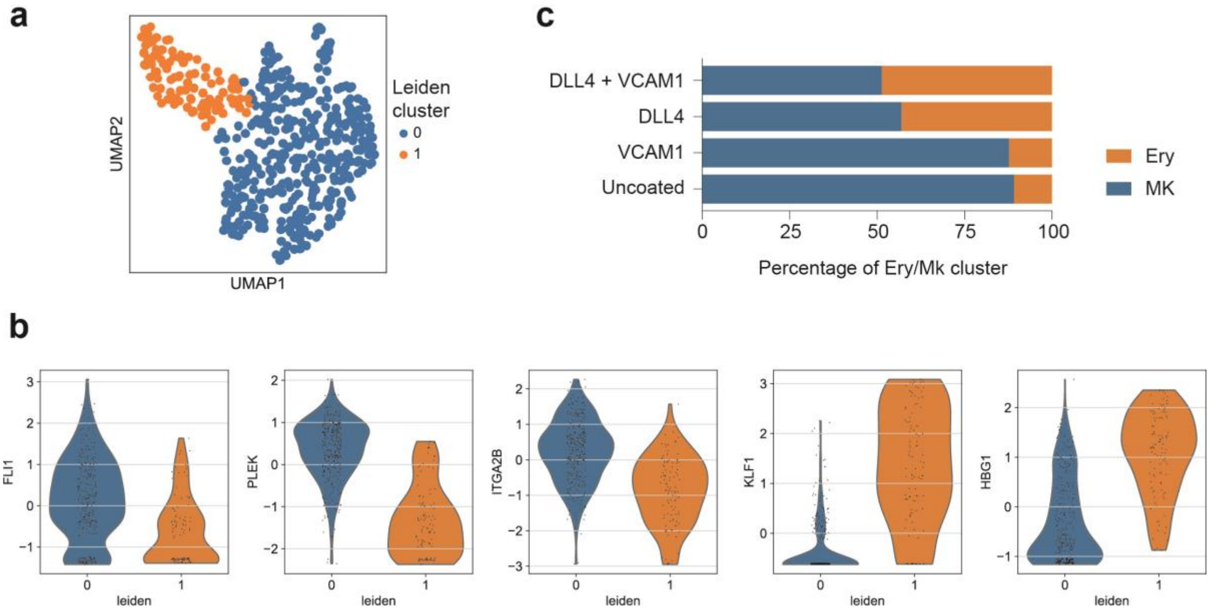


**Supplementary Figure 3: Addition of DLL4 during EHT increases the ratio of erythroid to megakaryocyte progenitors. a.)** Unsupervised sub-clustering of the Ery/MK progenitors reveal two populations. **b.)** Violin plots of known marker genes demonstrate that cluster 0 displays increased expression of the megakaryocyte-associated genes FLI1, PLEK and ITGA2B while cluster 1 expresses increased levels of the erythroid-associated genes KLF1 and HBG1. **c.)** Clusters from (a) were annotated based on marker gene expression in (b) and the relative abundance of erythroid (Ery) and megakaryocyte (MK) progenitor sub-clusters within the Ery/MK population are plotted for each coating condition.

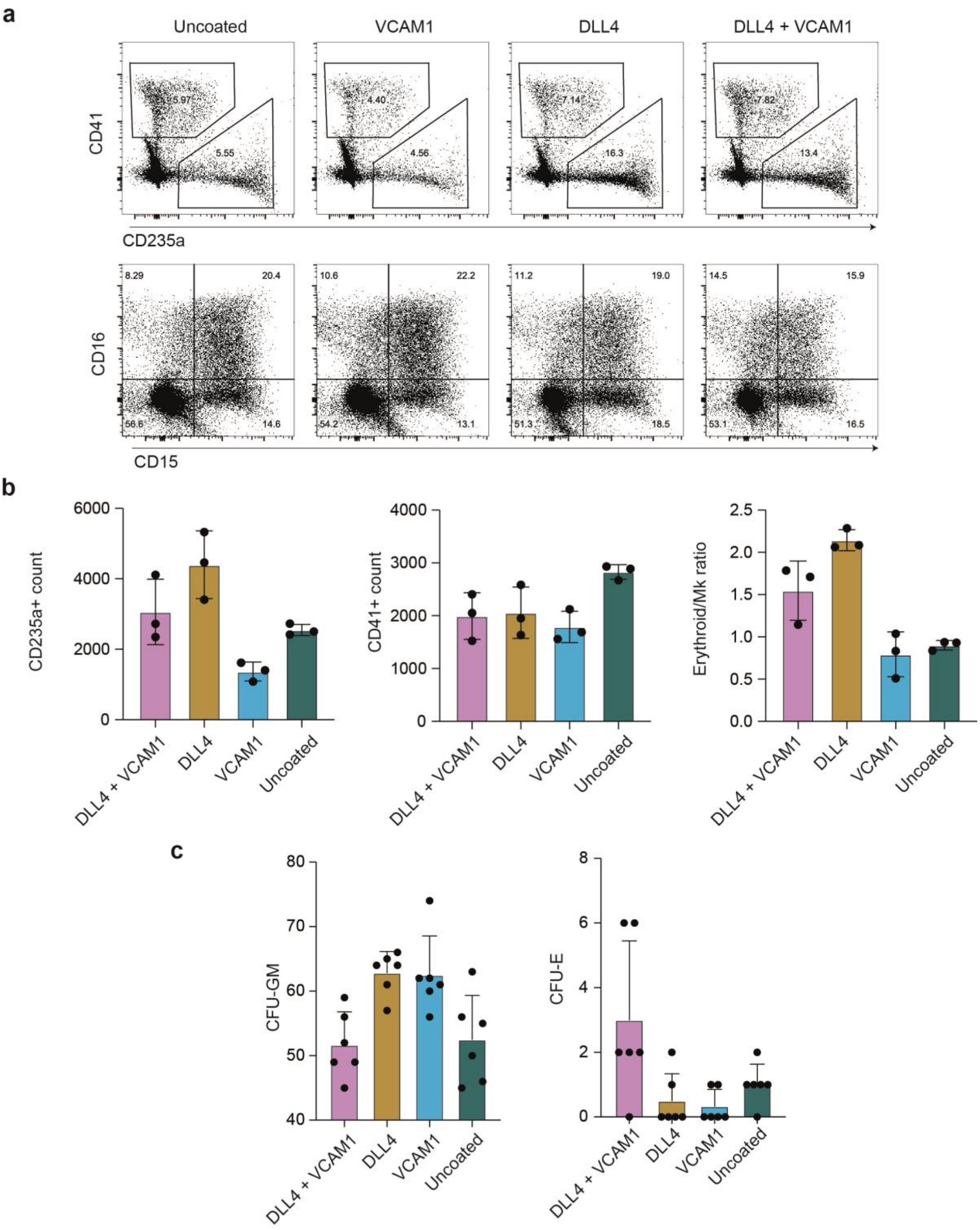


**Supplementary Figure 4: Addition of DLL4 during EHT alters erythroid, megakaryocyte and neutrophil output. a.)** Representative flow cytometry analysis of extended liquid cultures to mature non-lymphoid hematopoietic progenitors. **b.)** Yields and ratios of CD235+ erythroid cells and CD41+ megakaryocytes determined based on flow cytometry shown in (a). Yields are per well (n = 3, mean +/- s.d). Each well was initially seeded with 2500 PSC-derived cells harvested on day 5 of EHT. **c.)** Quantification of colony forming assays of cells harvested at day 5 of EHT from each coating condition (n=6, mean +s.d).

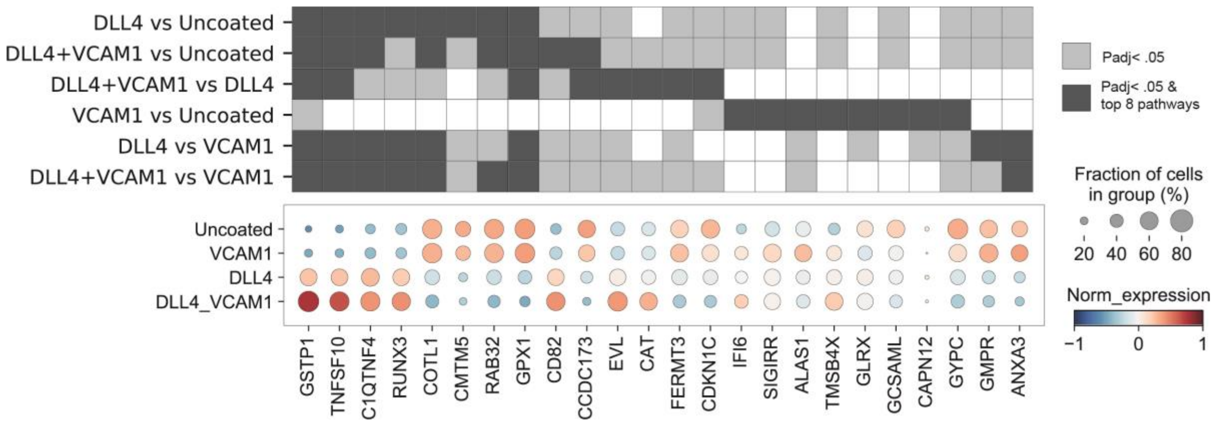


**Supplementary Figure 5: Unbiased exploration of the impact of DLL4 and VCAM1 during EHT on gene expression in PSC derived HSC/MPP.** Differential gene expression within HSC/MPP across coating conditions. Pairwise comparisons were performed between each coating condition and all others and the top 8 differentially expressed genes from each comparison by Z-score are depicted (t-test with overestimated variance, Benjamini-Hochberg corrected p-value correction) where a given gene was amongst the top 8 most differentially expressed.

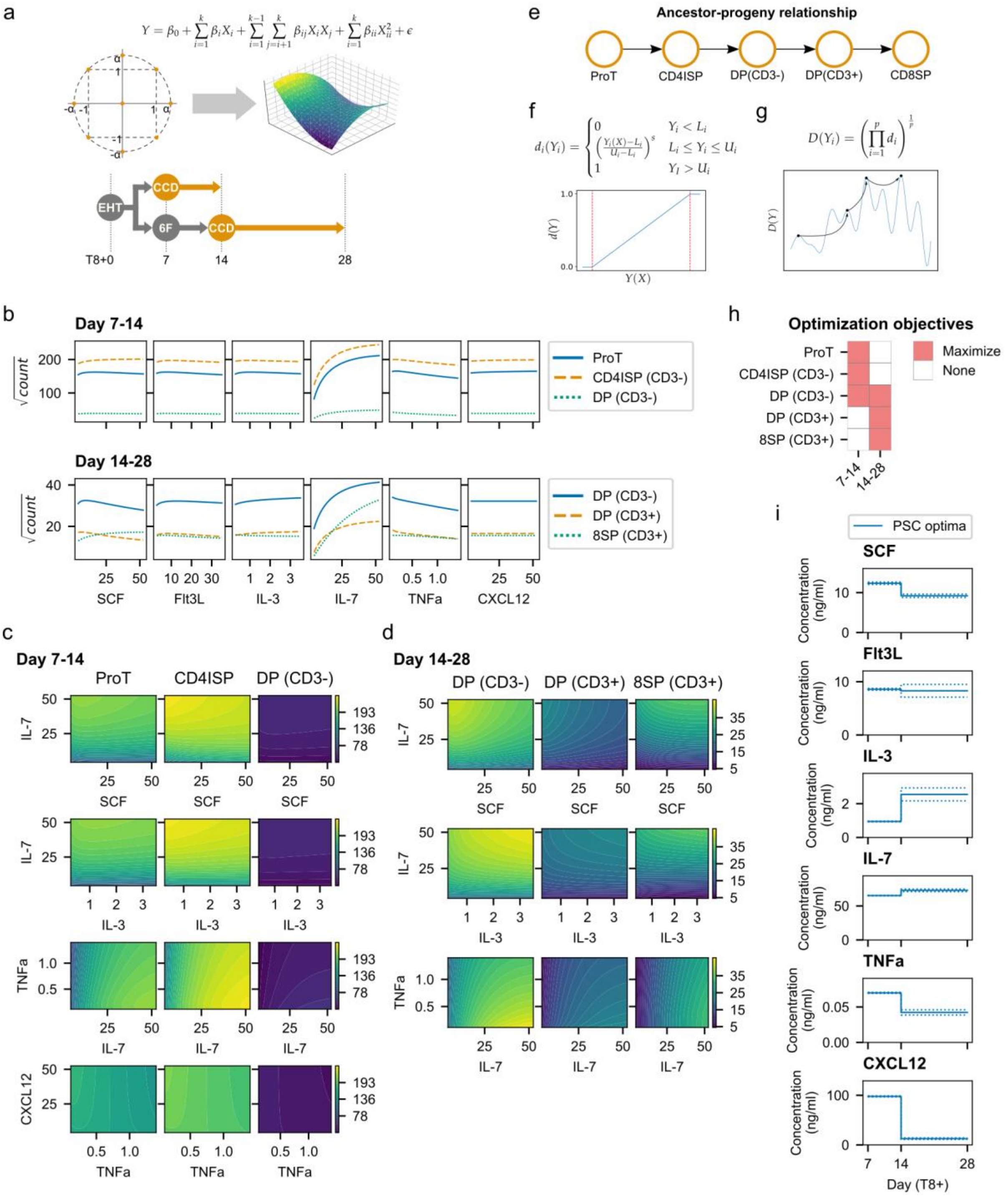


**Supplemental Figure 6. Modelling cytokine dose responses and optimization throughout T-cell development**. **a.)** A 6-factor orthogonal central composite design (CCD) experiment was performed at two stages of T-cell differentiation (T8+7-14 and T8+14-28). A polynomial equation fit using least-squares regression was used to model the dose response for each population of interest. **b.)** Predicted dose response for each population measured and for each cytokine. **c.)** Significant two-factor interactions between cytokines during day 7-14. **d.)** Significant two-factor interactions between cytokines during day 14-28. **e.)** In order to optimize cytokine concentrations to generate CD8+ T-cells, an ancestor-progeny relationship was assumed, where increasing the number of early phenotypes (ie. proT-cells and CD4ISP) would lead to larger numbers of DP and CD8SP T-cells later. **(f)** For each population of interest $i$, a desirability function $d(Y_i)$ was defined which scales the output of each polynomial model $Y_i(X)$ between [0,1]. Cytokine concentrations $X$ that increased $Y_i$ result in a desirability closer to 1 (more desirable) whereas those that decrease $Y_i$ are closer to 0 (less desirable). **g.)** The desirability function for each population was combined using the geometric mean to provide an overall desirability score $D$ which was optimized using the single objective basin-hopping algorithm. **h.)** Objective for each population of interest. Day 7-14 focused on early phenotypes while day 14-28 focused on more mature phenotypes once they were present in culture. **i.)** Predicted optimal cytokines for each stage. 25 random cytokine concentrations were used to initialize the basin-hopping algorithm and the top 5 most desirable solutions were kept. The solid line represents the mean of the top 5 solutions while the dotted line is the standard deviation. A larger standard deviation indicates that optimal solutions were less sensitive to that particular cytokine.

### Supplementary Tables

**Supplementary Table 1**. Cytokine concentrations tested in dose response experiments (ng/ml).

| Cytokine | -2.366 | -1 | 0 | 1 | 2.366 |
|---|---|---|---|---|---|
| SCF | 0.77 | 4.29 | 15 | 52.5 | 290.64 |
| Flt3L | 0.52 | 2.86 | 10 | 35 | 193.76 |
| IL-3 | 0.05 | 0.29 | 1 | 3.5 | 19.38 |
| IL-7 | 0.77 | 4.29 | 15 | 52.5 | 290.64 |
| TNFa | 0.02 | 0.11 | 0.4 | 1.4 | 7.75 |
| CXCL12 | 0.77 | 4.29 | 15 | 52.5 | 290.64 |

**Supplementary Table 2**. Regression coefficient estimates and statistics for sqrt[proT] during T8+7-14.

| Term | Estimate | Std Error | t Ratio | Prob>\|t\| |
|---|---|---|---|---|
| Intercept | 162.32027 | 4.08775 | 39.71 | <.0001 |
| SCF | 1.1753401 | 2.161302 | 0.54 | 0.59 |
| Flt3L | -2.297715 | 2.161302 | -1.06 | 0.295 |
| IL-3 | 0.3934496 | 2.161302 | 0.18 | 0.8566 |
| IL-7 | 73.429514 | 3.19335 | 22.99 | <.0001 |
| TNFa | -10.05826 | 2.161302 | -4.65 | <.0001 |
| CXCL12 | 2.6194807 | 2.161302 | 1.21 | 0.2336 |
| SCF*IL-7 | -5.546098 | 2.511207 | -2.21 | 0.0338 |
| IL-3*IL-7 | 1.0284591 | 2.511207 | 0.41 | 0.6846 |
| IL-7*TNFa | 0.4123762 | 2.511207 | 0.16 | 0.8705 |
| TNFa*CXCL12 | -5.542122 | 2.511207 | -2.21 | 0.034 |
| SCF*SCF | -6.541473 | 1.84083 | -3.55 | 0.0011 |
| Flt3L*Flt3L | -5.618722 | 1.84083 | -3.05 | 0.0043 |
| IL-3*IL-3 | -5.199209 | 1.84083 | -2.82 | 0.0078 |
| IL-7*IL-7 | -15.27823 | 1.84083 | -8.3 | <.0001 |
| TNFa*TNFa | -7.987426 | 1.84083 | -4.34 | 0.0001 |
| Block[1] | -11.28556 | 2.675735 | -4.22 | 0.0002 |
| Block[2] | -9.081582 | 2.71645 | -3.34 | 0.002 |
| IL-7*IL-7*IL-7 | -8.902384 | 1.072155 | -8.3 | <.0001 |

**Supplementary Table 3**. Regression coefficient estimates and statistics for sqrt[CD4ISP] during T8+7-14.

| Term | Estimate | Std Error | t Ratio | Prob>\|t\| |
|---|---|---|---|---|
| Intercept | 197.23105 | 3.449441 | 57.18 | <.0001 |
| SCF | 6.9739421 | 1.823812 | 3.82 | 0.0005 |
| Flt3L | -0.126888 | 1.823812 | -0.07 | 0.9449 |
| IL-3 | -0.552305 | 1.823812 | -0.3 | 0.7638 |
| IL-7 | 69.118812 | 2.694703 | 25.65 | <.0001 |
| TNFa | -8.101824 | 1.823812 | -4.44 | <.0001 |
| CXCL12 | 1.9329364 | 1.823812 | 1.06 | 0.2965 |
| SCF*IL-7 | -16.24081 | 2.119078 | -7.66 | <.0001 |
| IL-3*IL-7 | -4.824599 | 2.119078 | -2.28 | 0.029 |
| IL-7*TNFa | 1.2495945 | 2.119078 | 0.59 | 0.5592 |
| TNFa*CXCL12 | -4.13415 | 2.119078 | -1.95 | 0.0591 |
| SCF*SCF | -2.913235 | 1.553381 | -1.88 | 0.0691 |
| Flt3L*Flt3L | -5.529862 | 1.553381 | -3.56 | 0.0011 |
| IL-3*IL-3 | -2.913311 | 1.553381 | -1.88 | 0.0691 |
| IL-7*IL-7 | -13.1999 | 1.553381 | -8.5 | <.0001 |
| TNFa*TNFa | -5.861526 | 1.553381 | -3.77 | 0.0006 |
| Block[1] | -9.370574 | 2.257915 | -4.15 | 0.0002 |
| Block[2] | -6.356163 | 2.292272 | -2.77 | 0.0088 |
| IL-7*IL-7*IL-7 | -8.829736 | 0.904736 | -9.76 | <.0001 |

**Supplementary Table 4**. Regression coefficient estimates and statistics for sqrt[DP(CD3-)] during T8+7-14.

| Term | Estimate | Std Error | t Ratio | Prob>\|t\| |
|---|---|---|---|---|
| Intercept | 39.03312 | 1.007009 | 38.76 | <.0001 |
| SCF | 0.212637 | 0.532433 | 0.4 | 0.692 |
| Flt3L | -0.549534 | 0.532433 | -1.03 | 0.3091 |
| IL-3 | -0.274043 | 0.532433 | -0.51 | 0.61 |
| IL-7 | 14.123472 | 0.786675 | 17.95 | <.0001 |
| TNFa | -4.646284 | 0.532433 | -8.73 | <.0001 |
| CXCL12 | -0.044756 | 0.532433 | -0.08 | 0.9335 |
| SCF*IL-7 | -0.312975 | 0.618631 | -0.51 | 0.6161 |
| IL-3*IL-7 | -0.422433 | 0.618631 | -0.68 | 0.4992 |
| IL-7*TNFa | -2.074962 | 0.618631 | -3.35 | 0.0019 |
| TNFa*CXCL12 | -1.771969 | 0.618631 | -2.86 | 0.007 |
| SCF*SCF | -1.272841 | 0.453485 | -2.81 | 0.0081 |
| Flt3L*Flt3L | -1.444519 | 0.453485 | -3.19 | 0.003 |
| IL-3*IL-3 | -0.457725 | 0.453485 | -1.01 | 0.3197 |
| IL-7*IL-7 | -2.377885 | 0.453485 | -5.24 | <.0001 |
| TNFa*TNFa | -1.355425 | 0.453485 | -2.99 | 0.0051 |
| Block[1] | -2.890302 | 0.659162 | -4.38 | 0.0001 |
| Block[2] | -0.28152 | 0.669192 | -0.42 | 0.6766 |
| IL-7*IL-7*IL-7 | -1.876824 | 0.264123 | -7.11 | <.0001 |

**Supplementary Table 5**. Regression coefficient estimates and statistics for sqrt[DP(CD3-)] during T8+14-28.

| Term | Estimate | Std Error | t Ratio | Prob>\|t\| |
|---|---|---|---|---|
| Intercept | 32.174743 | 1.276881 | 25.2 | <.0001 |
| SCF | -1.939846 | 1.127087 | -1.72 | 0.0936 |
| Flt3L | 0.5175565 | 0.760114 | 0.68 | 0.5002 |
| IL-3 | 1.5683656 | 0.760114 | 2.06 | 0.0461 |
| IL-7 | 12.935004 | 1.127087 | 11.48 | <.0001 |
| TNFa | -2.438299 | 1.127087 | -2.16 | 0.037 |
| SCF*IL-7 | -1.920256 | 0.88782 | -2.16 | 0.0371 |
| IL-3*IL-7 | 1.9574174 | 0.88782 | 2.2 | 0.0338 |
| IL-7*TNFa | -2.504479 | 0.88782 | -2.82 | 0.0077 |
| SCF*SCF | -2.756734 | 0.636598 | -4.33 | 0.0001 |
| Flt3L*Flt3L | -1.398024 | 0.636598 | -2.2 | 0.0344 |
| IL-7*IL-7 | -2.045512 | 0.636598 | -3.21 | 0.0027 |
| TNFa*TNFa | -1.294971 | 0.636598 | -2.03 | 0.0491 |
| SCF*SCF*SCF | 0.3599793 | 0.373356 | 0.96 | 0.3412 |
| IL-7*IL-7*IL-7 | -1.76639 | 0.373356 | -4.73 | <.0001 |
| TNFa*TNFa*TNFa | -0.716608 | 0.373356 | -1.92 | 0.0627 |

**Supplementary Table 6**. Regression coefficient estimates and statistics for sqrt[DP(CD3+)] during T8+14-28.

| Term | Estimate | Std Error | t Ratio | Prob>\|t\| |
|---|---|---|---|---|
| Intercept | 16.533849 | 0.819832 | 20.17 | <.0001 |
| SCF | -2.023788 | 0.723655 | -2.8 | 0.0081 |
| Flt3L | -0.365552 | 0.488037 | -0.75 | 0.4586 |
| IL-3 | 0.9113174 | 0.488037 | 1.87 | 0.0698 |
| IL-7 | 8.5043609 | 0.723655 | 11.75 | <.0001 |
| TNFa | -1.488186 | 0.723655 | -2.06 | 0.0468 |
| SCF*IL-7 | -1.923721 | 0.570032 | -3.37 | 0.0017 |
| IL-3*IL-7 | 1.1479938 | 0.570032 | 2.01 | 0.0513 |
| IL-7*TNFa | -0.908999 | 0.570032 | -1.59 | 0.1193 |
| SCF*SCF | -1.377391 | 0.408733 | -3.37 | 0.0018 |
| Flt3L*Flt3L | -1.109826 | 0.408733 | -2.72 | 0.01 |
| IL-7*IL-7 | -1.641637 | 0.408733 | -4.02 | 0.0003 |
| TNFa*TNFa | -0.381667 | 0.408733 | -0.93 | 0.3565 |
| SCF*SCF*SCF | 0.1718311 | 0.239716 | 0.72 | 0.478 |
| IL-7*IL-7*IL-7 | -1.008564 | 0.239716 | -4.21 | 0.0002 |
| TNFa*TNFa*TNFa | -0.657095 | 0.239716 | -2.74 | 0.0094 |

**Supplementary Table 7**. Regression coefficient estimates and statistics for log[8SP + 1] during T8+14-28.

| Term | Estimate | Std Error | t Ratio | Prob>\|t\| |
|---|---|---|---|---|
| Intercept | 5.4937311 | 0.113162 | 48.55 | <.0001 |
| SCF | 0.3277836 | 0.099887 | 3.28 | 0.0023 |
| Flt3L | -0.075585 | 0.067364 | -1.12 | 0.2691 |
| IL-3 | -0.050463 | 0.067364 | -0.75 | 0.4585 |
| IL-7 | 1.891356 | 0.099887 | 18.94 | <.0001 |
| TNFa | -0.060797 | 0.099887 | -0.61 | 0.5465 |
| SCF*IL-7 | -0.077879 | 0.078682 | -0.99 | 0.3287 |
| IL-3*IL-7 | 0.2352338 | 0.078682 | 2.99 | 0.0049 |
| IL-7*TNFa | -0.001178 | 0.078682 | -0.01 | 0.9881 |
| SCF*SCF | -0.082605 | 0.056418 | -1.46 | 0.1516 |
| Flt3L*Flt3L | -0.097429 | 0.056418 | -1.73 | 0.0925 |
| IL-7*IL-7 | -0.244079 | 0.056418 | -4.33 | 0.0001 |
| TNFa*TNFa | -0.104099 | 0.056418 | -1.85 | 0.073 |
| SCF*SCF*SCF | -0.047102 | 0.033088 | -1.42 | 0.163 |
| IL-7*IL-7*IL-7 | -0.162448 | 0.033088 | -4.91 | <.0001 |
| TNFa*TNFa*TNFa | -0.056402 | 0.033088 | -1.7 | 0.0967 |
